# A Recurrent Neural Network Based Method for Genotype Imputation on Phased Genotype Data

**DOI:** 10.1101/821504

**Authors:** Kaname Kojima, Shu Tadaka, Fumiki Katsuoka, Gen Tamiya, Masayuki Yamamoto, Kengo Kinoshita

## Abstract

Genotype imputation estimates genotypes of unobserved variants from genotype data of other observed variants, and such estimation is enabled using haplotype data of a large number of other individuals. Although existing imputation methods explicitly use haplotype data, the accessibility of haplotype data is often limited because the agreement is necessary from donors of genome data. We propose a new imputation method that uses bidirectional recurrent neural network, and haplotype data of a large number of individuals are encoded as its model parameters through the training step, which can be shared publicly due to the difficulty in restoring genotype data at the individual-level. In the performance evaluation using the phased genotype data in the 1000 Genomes Project, the imputation accuracy of the proposed method in *R*^2^ is comparative with existing methods for variants with MAF ≥ 0.05 and is slightly worse than those of the existing methods for variants with MAF < 0.05. In a scenario of limited availability of haplotype data to the existing methods, the accuracy of the proposed method is higher than those of the existing methods at least for variants with MAF ≥ 0.005. Python code of our implementation for imputation is available at https://github.com/kanamekojima/rnnimp/.

## 1 Introduction

The development of high-throughput sequencing technologies enables the construction of genotype data with base-level resolution for more than one thousand individuals. Such large-scale, high-resolution genotype data is often called reference panel, and one of applications of the reference panel is genotype imputation. In SNP array, genotype data can be obtained with lower cost than sequencing, while available genotypes are limited to designed markers. The current imputation methods take phased genotypes obtained by SNP array or other genotyping technologies as input genotype data, and estimate haplotypes matching to the input genotype data from the recombination of haplotypes in the reference panel based on Li and Stephens model [Li and Stephens, 2003]. Genotypes of unobserved variants are then obtained from the estimated haplotypes.

Although existing imputation methods such as Impute2 [Howie *et al*., 2009], Minimac3 [Das *et al*., 2016], and Beagle5 [Browning *et al*., 2018] explicitly use haplotype data, the accessibility of haplotype data is often limited because the agreement with donors of genome data is necessary for public use. For example, there are 64,976 haplotypes in the Haplotype Reference Consortium [McCarthy *et al*., 2016] in total, but the number of publicly available haplotypes is 22,454, and about two third of the remaining haplotypes can only be used inside of the imputation server. Thus, the input genotype data must be sent to other research institutes for imputation using publicly unavailable reference panel, although the input genotype data often has some limitations for external use due to the informed consent policy. One solution for this issue is to encode the information of phased genotype data to summary statistics or model parameters from which the restoration of genotype data at the individual-level is difficult. Recent development of deep learning techniques including recurrent neural networks provides the improvement of accuracy in various fields such as image classification [Krizhevsky *et al*., 2012], image detection [He *et al*., 2017], natural language understanding [Sutskever *et al*., 2014], and speech and video data recognition [Ephrat *et al*., 2018], and hence the use of deep learning techniques is one of the promising strategies for devising imputation methods handling haplotype information as model parameters.

In this study, we propose a new imputation method based on bidirectional recurrent neural network (RNN) that takes haplotype of phased genotypes as input data and returns estimated alleles for unobserved variants. In the proposed method, haplotype information is parameterized as model parameters in the training step, and haplotype data itself is not explicitly used in the imputation. We considered binary vectors indicating alleles in the reference panel for the feature information of variants in input data, which are converted to the binary vectors to make input feature vectors of bidirectional RNN using kernel principal component analysis. The aim of this conversion was to reduce the size of feature vectors and to avoid the restoration of genotype data. For RNN cells, we considered long short-term memory (LSTM) [Hochreiter and Schmidhuber, 1997] and gated recurrent unit (GRU) [Cho *et al*., 2014] in the proposed method. We also propose a hybrid model obtained by combining two bidirectional RNN models trained for different minor allele frequency (MAF) ranges. Since it is difficult to restore genotype information at individual-level from model parameters, the haplotype information with the permission of statistical reuse can be publicly used. For performance evaluation, we used phased genotype data of 2,504 individuals in the 1000 Genomes Project [1000 Genomes Project Consortium *et al*., 2015]. The imputation accuracy of the proposed model in *R*^2^ is comparative with those of existing methods for variant sites of MAF ≥ 0.05. For variant sites of MAF < 0.05, the imputation accuracy of the proposed model in *R*^2^ is slightly worse than those of existing methods. We also considered a condition for limited availability of haplotype data to existing methods, and showed that the accuracy of the proposed method was higher than those of other existing methods, at least for variants with MAF ≥ 0.005.

## 2 Methods

Let *v*_*i*_ and *u*_*j*_ be the *i*th observed variant and *j*th unobserved variant, respectively. The order of variants is sorted with their genomic positions, and hence *p*(*v*_*i*_) ≤ *p*(*v*_*j*_) and *p*(*u_i_*) ≤ *p*(*u*_*j*_) are satisfied for *i* < *j*, where *p*(·) is a function returns the position of input variant. We divided a chromosome to regions according to the numbers of observed and unobserved variants as shown in Fig. 1. We limited the maximum numbers of observed and unobserved variants in each region to 100 and 1,000 in our experiment, respectively, and regions were extended up to the limit in the division process. Each divided region had flanking regions in the upstream and downstream directions, and only observed variants were considered in the flanking regions. The proposed method takes haplotype of observed variants and imputes unobserved variants for each region, and imputation results from the divided regions are concatenated for the final imputation result. In the following subsections, we describe the model structure of the proposed method for each region, extraction of input feature vectors for the model, and procedures for training the model in details.

**Figure 1:**
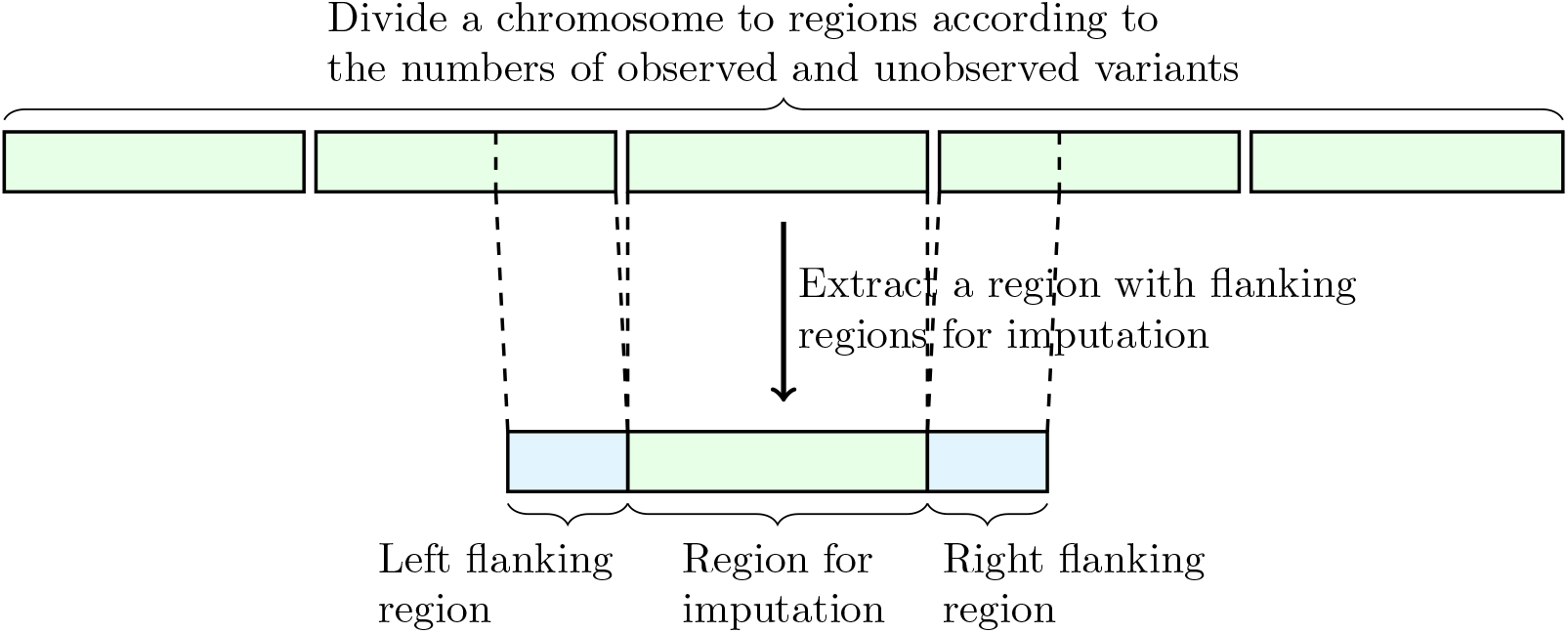
An illustration of division of a chromosome to regions according to the numbers of observed and unobserved variants for imputation.

### 2.1 Model structure

We assume that both observed and unobserved variants are biallelic; i.e., their alleles are represented by one and zero. Let *m* be the number of observed variants in a divided region. We also let *m*_*l*_ and *m*_*r*_ respectively be indices of left most and right most observed variants in the region without left and right flanking regions. We build bidirectional RNN on observed variants for each divided region as shown in Fig. 2. Forward RNN is built on observed variants 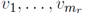, and observed variants 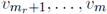 in the right flanking region are not included, since variants in the right flanking region are not required for imputing unobserved variants in the forward direction. Backward RNN is built on observed variants 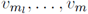, and similarly to the forward RNN, 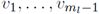 in the left flanking region are not included. RNN cells for each observed variant of forward and backward RNNs are stacked in the proposed model as shown in Fig. 3, and LSTM and GRU are considered for RNN cells. 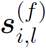 and 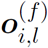 in Fig. 3 are the state and output vectors of the RNN cell for the *l*th layer on observed variant *v*_*i*_ in forward RNN, respectively. 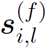 and 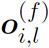 are obtained recursively for *i* ∈ {1,…, *m*_*r*_} in the following manner:

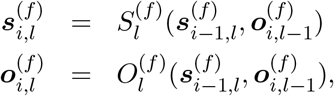

**Figure 2:**
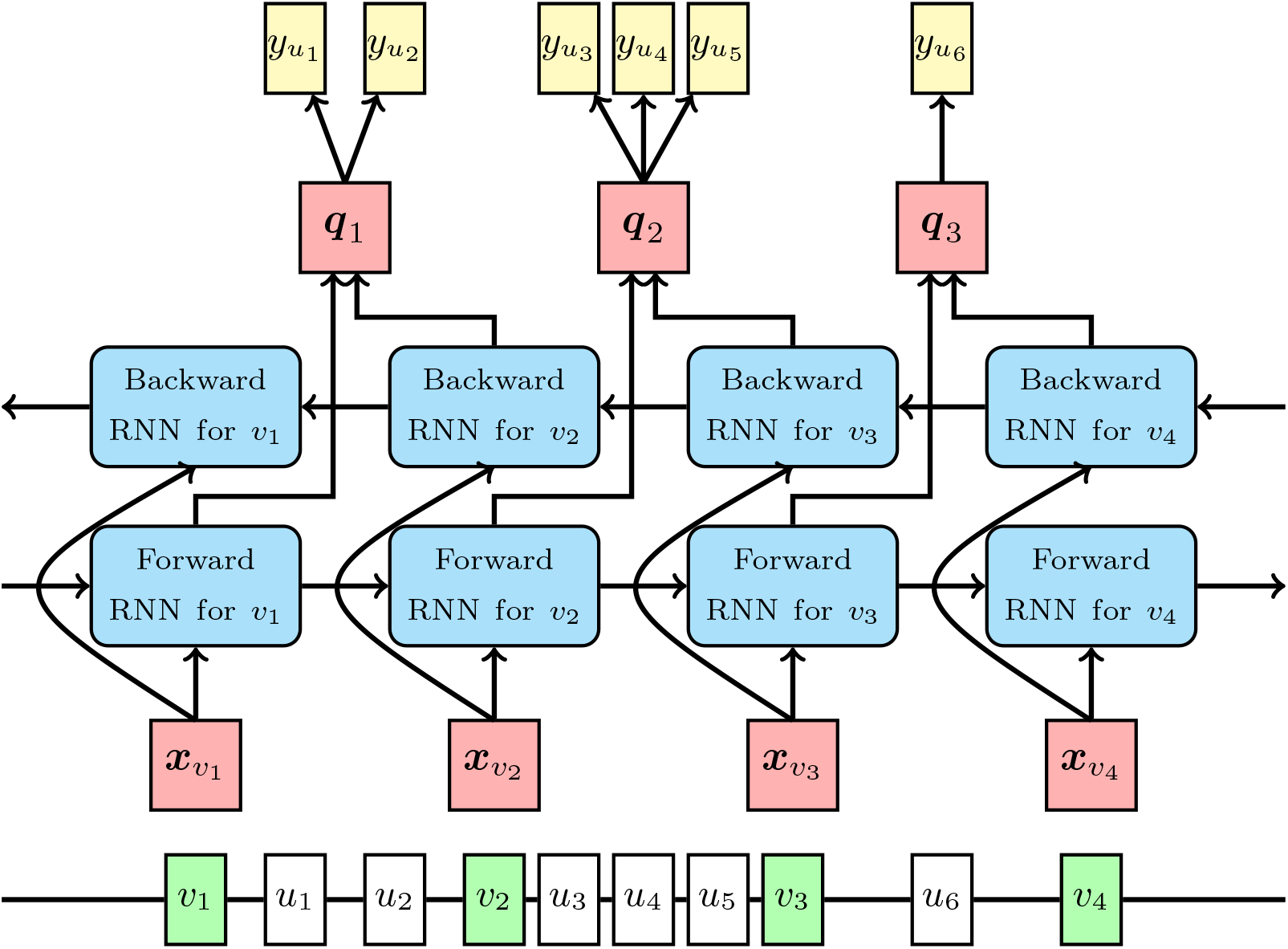
The overall model structure of the proposed method. The line in the bottom of the figure indicates a genome sequence where observed variants are in green square and unobserved variants are in white square. Forward and backward RNNs are built on observed variants. 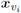 is the input feature vector of forward and backward RNNs for observed variant *v*_*i*_. ***q****i* is the vector from the concatenation of the output of the forward RNN for observed variant *v*_*i*_ and the output of the backward RNN for observed variant *v*_*i*+1_. 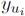 is a binary variable indicating the allele for unobserved variant *u*_*i*_.

**Figure 3:**
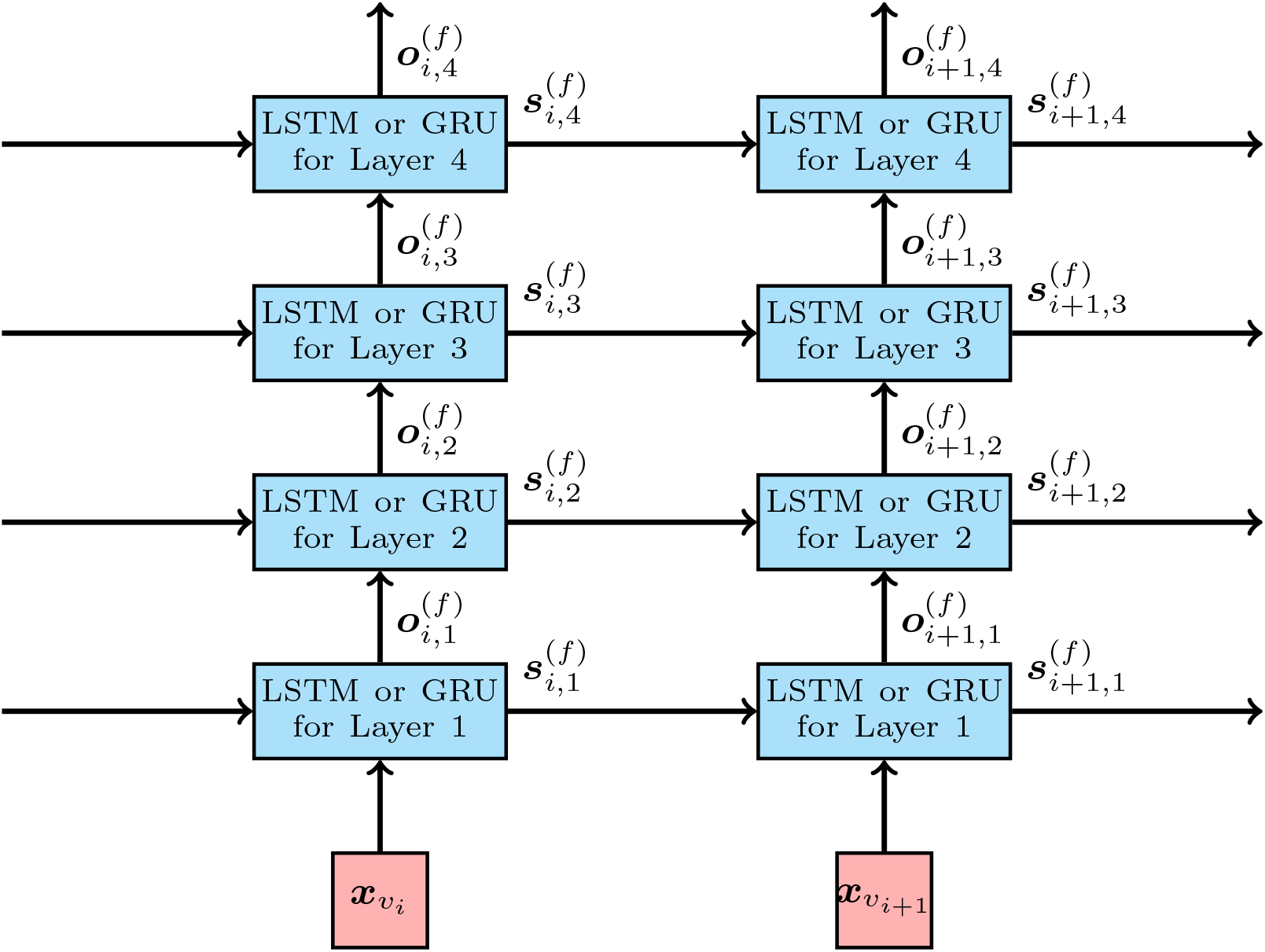
The RNN structure on each observed variant for the case of four stacked RNN cells in the forward RNN. 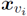 and 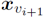 are respectively input feature vectors for observed variants *v*_*i*_ and *v*_*i*+1_. 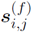 is the state of the RNN cell of the *j*th layer for observed variant *v*_*i*_ and used as the input of the state for the RNN cell of the *j*th layer for observed variant *v*_*i*+1_. The output of the RNN cell of the top layer, o_*i*,4_, is handled as the output of RNN for each observed variant.

where 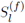 and 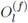 are functions that return the state and output vectors for the RNN cell of the *l*th layer, respectively. Note that the initial state 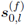 is set to **0**, and 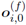 is set to input feature vector for the *i*th observed variant *x*_*v_i_*_. Details of input feature vectors for observed variants are described in the next subsection. For backward RNN, we used the corresponding notations to those used in forward RNN, and obtained

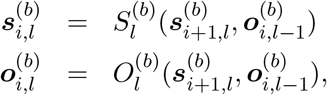

where *i* ∈ {*m*_*l*_,…, *m*}. Note that 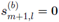 and 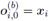.

Let ***q***_*i*_ be a vector given by concatenation of output vectors of forward and backward RNNs as shown in Fig. 2:

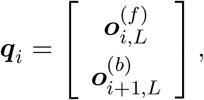

where *L* is the number of layers in the model. Let 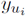 be a binary value representing the allele of unobserved variant *u*_*i*_. The probability of 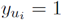 is estimated by the following softmax function:

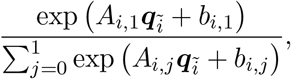

where *A*_*i*,*j*_ and *b*_*i*,*j*_ are parameters to be trained, and 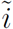 is the index that satisfies 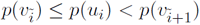; i.e., 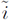 is the index for the closest observed variant to *u*_*i*_ in the upstream region. For the case of *p*(*u*_*i*_) < *p*(*v*_1_), which occurs in the left most divided region, we used 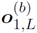 as 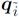. Similarly, we used 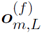 as 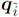 case of *p*(*u*_*i*_) ≥ *p*(*v*_*m*_).

For the loss function for the training of model parameters, we considered the sum of weighted cross entropies over the unobserved variants as follows:

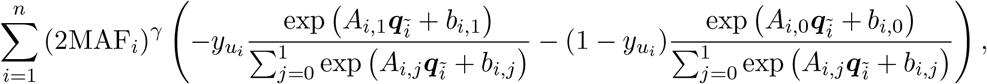

where *n* is the number of unobserved variants, MAF_*i*_ is the minor allele frequency of *u_i_* in the training data, and γ is a hyper-parameter to adjust the weights from MAF. The loss function with γ > 0 gives higher priority to higher MAF variants, while that with γ < 0 gives higher priority to lower MAF variants. Hereafter, we call the model trained with γ > 0 “higher MAF model” and that with γ < 0 “lower MAF model”. In order to take advantage of the two types of models for achieving higher accuracy in both high and low MAF variants, we propose a hybrid model obtained by the combination of the higher and lower MAF models. For the hybrid model, we consider the combination of logits of these two models of the *i* unobserved variant as ***r***_*i*_ defined as follows:

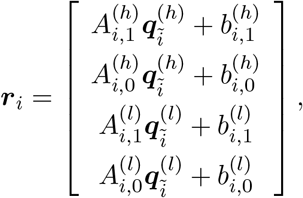

where superscripts (*h*) and (*l*) indicate variables and outputs of higher MAF model and lower MAF model, respectively, We then estimated the probability of 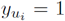 by the following softmax function for ***r***_*i*_:

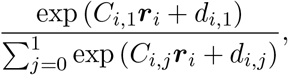

where *C*_*i*,*j*_ and *d*_*i*,*j*_ are parameters. After the learning of the parameters of the higher and lower MAF models, we trained *C*_*i*,*j*_ and *d*_*i*,*j*_ in the loss function by the sum of cross entropies as follows:

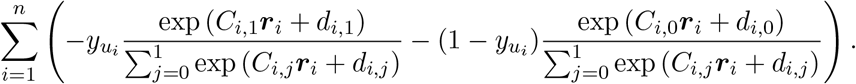

Note that the parameters of higher and lower MAF models were fixed for training *C*_*i*,*j*_ and *d*_*i*,*j*_.

### 2.2 Input feature vectors for observed variants in a reference panel

Let *B* be a binary matrix representing a reference panel, where the *i*th row and *j*th column element indicates the allele of the *i*th haplotype in the *j*th variant. We first consider the *j*th column vector of *B* as a feature vector for an allele indicated by one at the *j*th variant. For observed variant *v*, we denote the feature vector for the allele indicated by one as 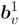. We also let 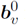 be the feature vector for an allele indicated by zero, in which the *i*th element takes one if the allele of the *i*th haplotype is indicated by zero, and zero otherwise. For example, let us consider the following allele pattern for a variant site with alleles ‘A’ and ‘T’ in a reference panel:

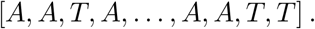

If ‘A’ and ‘T’ are respectively indicated by one and zero, the corresponding binary representation is given by

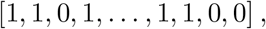

and feature vectors 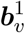 and 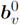 for allele ‘A’ and ‘T’ are given by [1, 1, 0, 1,…, 1, 1, 0, 0] and [0, 0, 1, 0,…, 0, 0, 1, 1], respectively. These feature vectors can be interpreted as a binary vector indicating which haplotypes have the input allele for the variant. However, there are two serious problems in the feature vectors; they are the explicit representation of a reference panel, and they are too big as an RNN input, since the number of individuals in a reference panel is usually more than 1,000.

Thus, we adopt kernel principal component analysis (PCA) [Schölkopf *et al*., 1998] as a dimensionality reduction technique for the feature vector in order to resolve the two issues at the same time. Since the correlation of 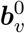 and 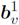 is minus one, we applied kernel PCA only to feature vectors for alleles indicated by one: 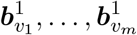, to manage the problem of PCA results caused by highly correlated variables. In order to obtain the reduced feature vector of 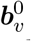, we projected 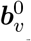 to the space from kernel PCA obtained for 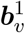. Given an original binary feature vector **b**, the *i*th element of its dimensionally reduced feature vector is given by

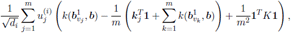

where *k*(·, ·) is a positive definite kernel, *K* is Gram matrix, ***k***_*i*_ is the *i*th column vector of *K*, *d*_*i*_ is the *i*th largest eigenvalue of the centered Gram matrix 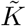, and 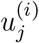 is the *j*th element of the corresponding eigenvector of *d*_*i*_. Details of the derivation for the above equation are in Section S1.

### 2.3 Training of the proposed model

We used Adam optimizer [Kingma and Ba, 2015] to train parameters of the proposed model. In order to avoid overfitting of parameters, we considered averaged cross entropy loss and *R*^2^ value in the validation data as early stopping criteria. Note that *R*^2^ value was obtained as the squared correlation of true genotype count and allele dosage. In the practical trials, we found that averaged *R*^2^ value in the validation data was suitable for lower MAF variants, while the cross entropy loss for the validation data was suitable for the higher MAF variants. Thus, we used the cross entropy loss for the validation data as the early stopping criterion for training higher MAF model, and *R*^2^ value in the validation data for lower MAF model and hybrid model in the following results. In the training step, we reduced the learning rate if the early stopping criterion was not updated in the specified number of iterations, which we called the learning rate updating interval. Training stops if the learning rate gets less than the minimum learning rate or the iteration count reaches the maximum iteration count. Details of the training step are summarized as follows:

1. Set iteration count *i* to 1 and set the best value for the early stopping criterion ĉ to null.
2. Set learning rate *lr* and learning rate updating interval *li* to some initial values.
3. If *i* is larger than the maximum iteration count, finish training.
4. Update model parameters by Adam optimizer with learning rate *lr* for randomly selected batch data.
5. If *i* is divisible by validation interval *vi*, compute the following procedures:

a. Calculate the current value for the early stopping criterion *c*.
b. If ĉ is null or *c* is better than *ĉ*, set *ĉ* to *c*, save the current parameters, and set the last parameter saving step *i_s_* to *i*.
c. If *i* − *i*_*s*_ is larger than learning rate updating interval *li*:

i. Divide learning rate *lr* by two.
ii. If learning rate *lr* is less than the minimum learning rate *lr*_min_, finish training.
iii. Divide learning rate updating interval *li* by two and round it down to an integer.
iv. If learning rate updating interval *li* is less than the minimum learning rate updating interval *li*_min_, set *li* to *li*_min_.
v. Set the last parameter saving step *i*_*s*_ to *i*.
vi. Restore parameters to the previously saved parameters.
6. Increment *i* and go back to Step 2.

Since the local search in less space is expected for the smaller learning rate in the above procedures, we reduced the learning rate updating interval along with the learning rate. In our experiments, we set initial learning rate to 0.0001, the minimum learning rate *lr* _min_ to 10^−7^, initial learning rate updating interval to 5,000, the minimum learning rate updating interval *li*_min_ to 100, validation interval *vi* to 10, and the maximum iteration count to 100,000.

## 3 Results and Discussion

We used phased genotype data of 2,504 individuals for chromosome 22 from the phase 3 data of the 1000 Genomes Project [1000 Genomes Project Consortium *et al*., 2015]. We randomly selected 100 individuals for test data and evaluated the imputation performance for the test data by using the phased genotype data of the remaining 2,404 individuals as the reference panel. In the test data, we extracted genotype data for designed markers in SNP array and impute genotypes for variants from the extracted genotype data by using a reference panel. We first examined the imputation accuracy of the proposed method for the following the number of layers, the number of hidden units, and RNN cell types:

- RNN cell type: LSTM or GRU
- The number of layers: 2 or 4
- The number of hidden units: 20 or 40

The proposed method was implemented in Python 3 and TensorFlow (https://www.tensorflow.org/) was used for the implementation of RNN. We extracted genotypes for designed markers in Infinium Omni2.5-8 BeadChip, which we have called Omni2.5 hereafter, in the test data. The number of markers designed in Omni2.5 in chromosome 22 is 31,325, and 1,078,043 variants in the reference panel that were not in the designed markers in Omni2.5 were evaluated for the imputation accuracy. It should be noted that we filtered out variants with MAF < 0.005 for imputation, because the rare variants are not usually used for later analyses, although high computation cost is required for imputing all the variant in the proposed method. As a positive definite kernel for feature extraction, we used the following homogeneous dot-product kernel [Mairal, 2016]:

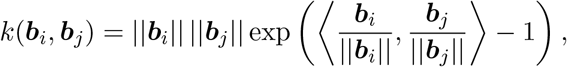

where ∥·∥ indicates L2 norm of input vector and 〈·, ·〉 indicates the inner product of two input vectors. For the loss function of the proposed method, γ was set to 0.75. We compared averaged *R*^2^ values in validation data for cases of using input feature vectors with top 5, 10, and 20 principal component scores as shown in Table 1. In the comparison, the proposed model with GRU, 4 layers, and 40 hidden units was used, and the average *R*^2^ value for the case of top 10 principal component scores was higher than those of other cases, although the size of input feature vectors was not sensitive to the imputation accuracy. Based on the comparison, we used top 10 principal component scores for the input feature vector in the following experiments. Table 2 shows averaged *R*^2^ value in the validation data for each setting. Fig. 4(a) shows the comparison of *R*^2^ values for settings with minimum or maximum averaged *R*^2^ value for LSTM and GRU. The proposed model with GRU, 4 layers, and 40 hidden units gave the highest averaged *R*^2^ value in the validation data among the settings, and the comparison of averaged *R*^2^ values in the validation data was consistent with the results in the test data. In order to see the effectiveness of the hybrid model in the proposed method, we compared the *R*^2^ values of results of hybrid model, higher MAF model, and lower MAF model. From the comparison of *R*^2^ values in Fig. 4(b), the hybrid model was comparable with higher MAF model in higher MAF range. In lower MAF range, the hybrid model was comparable with lower MAF model and better than the model for higher MAF variants. Hence, the hybrid model was effective over the entire MAF range compared with the higher and lower MAF models.

**Table 1:** Averaged *R*^2^ values in the validation data for input feature vectors with size of 5, 10, and 20. Size of Input Feature Vector 5, 10, and 20.

| Size of Input Feature Vector | 5 | 10 | 20 |
| --- | --- | --- | --- |
| $R^2$ | 0.8707 | 0.8708 | 0.8705 |

**Table 2:** Averaged *R*^2^ values in the validation data for several settings.

| RNN cell | No. of<br>Layers | No. of<br>Hidden Units | $R^2$ |
| --- | --- | --- | --- |
| LSTM | 2 | 20 | 0.8671 |
|  | 2 | 40 | 0.8701 |
|  | 4 | 20 | 0.8667 |
|  | 4 | 40 | 0.8690 |
| GRU | 2 | 20 | 0.8673 |
|  | 2 | 40 | 0.8702 |
|  | 4 | 20 | 0.8674 |
|  | 4 | 40 | 0.8708 |

**Figure 4:**
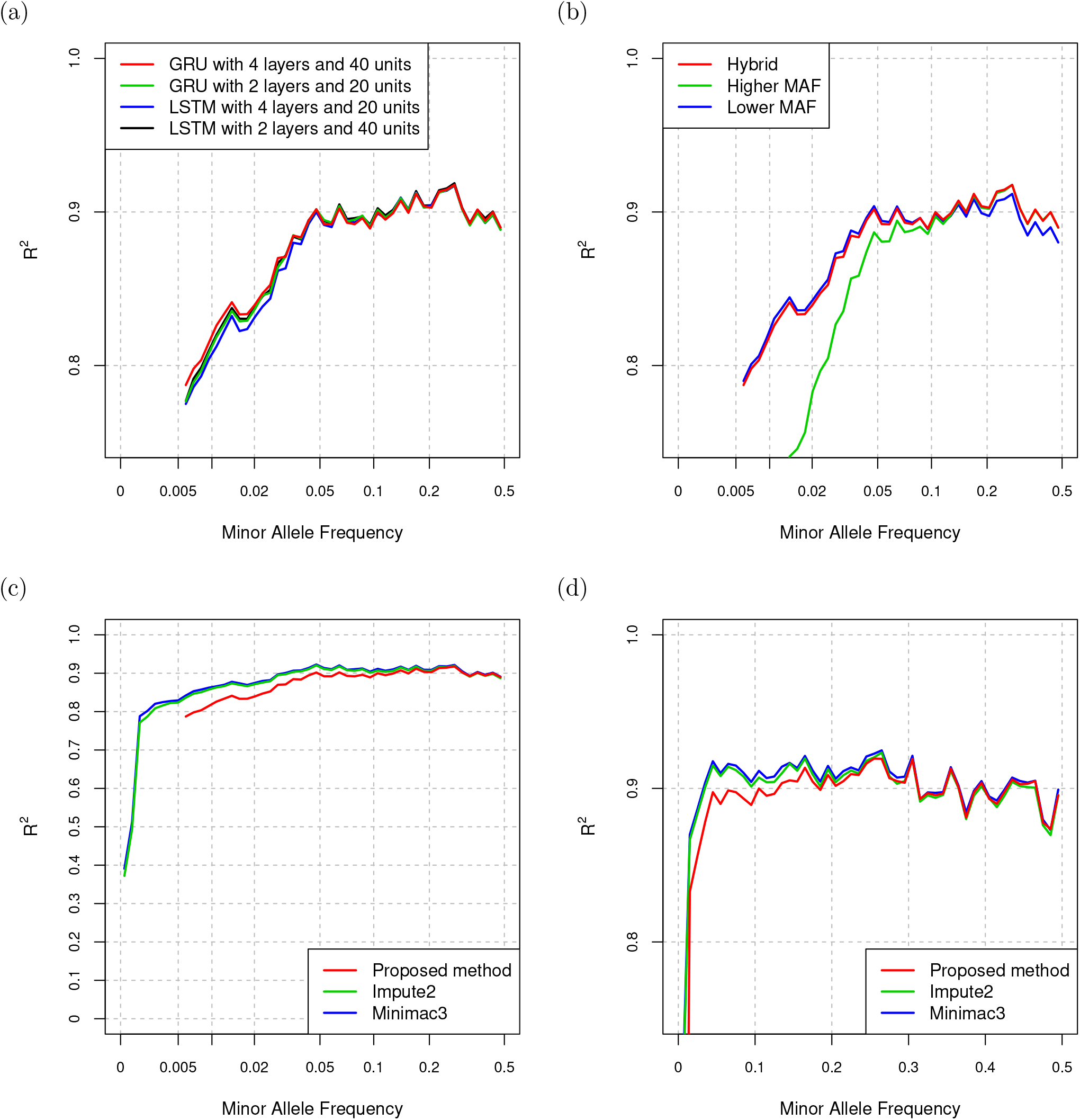
(a) Comparison of *R*^2^ values for the proposed method with several settings. (b) Comparison of *R*^2^ values for the proposed method with hybrid model, higher MAF model, and lower MAF model with the setting of GRU, 4 layers, and 40 hidden units. (c) Comparison of *R*^2^ values for the proposed method, Impute2, and Minimac3. (d) Comparison of *R*^2^ values for the proposed method, Impute2, and Minimac3 in linear MAF scale and with zoom into higher *R*^2^ value.

We selected Impute2 and Minimac3 as the representatives of existing imputation methods, and compared the imputation performance of the proposed method and these methods. For the proposed method, we used hybrid model with the setting of GRU, 4 layers, and 40 hidden units. Fig. 4(c) shows the comparison of *R*^2^ values. The imputation accuracy of the proposed method was worse than those of existing methods for MAF < 0.05 in *R*^2^. Fig. 4(d) shows the comparison of *R*^2^ values in linear MAF scale and with zoom into higher *R*^2^ value. For the higher MAF, the accuracy of the proposed method was comparable with those of existing methods. Especially for MAF ¿ 0.3, the proposed method was slightly better than Impute2. We also compared the running time of the proposed method and the existing methods in Table 3. All experiments were performed on Intel Xeon Silver 4116 CPU (2.10GHz). Although the proposed method required approximately two times as much running time as Impute2 for the imputation using trained parameters, the running time was feasible for practical use. Since running time of the proposed method is highly dependent on TensorFlow, the reduction of running time is expected along with the development of TensorFlow.

**Table 3:** Running time of the proposed and existing methods

| Method | Running Time for Imputation | Running Time for Training |
| --- | --- | --- |
| Proposed | 28,570 [s] | 13,471,091 [s] |
| Impute2 | 13,998 [s] | NA |
| Minimac3 | 7,374 [s] | NA |

We considered the case that genotype data of some individuals are not publicly available, but can be used for training the model of proposed method. In order to evaluate the imputation performance for the case, we randomly selected 100 EAS individuals of 504 EAS individuals for the EAS test data, and prepared two types of reference panels: a reference panel comprised of the remaining 2,404 individuals, and a reference panel with no EAS individuals comprised of 2,000 individuals. We used the former reference panel for training the proposed model, and used the latter panel for the imputation with the existing methods. In the evaluation, we considered Omni2.5 as the SNP array in the test data. For the proposed method, we also used hybrid model with the setting of GRU, 4 layers, and 40 hidden units. Fig. 5 shows the comparison of *R*^2^ values for the test data in scaled MAF ranges. At least for MAF ≥ 0.005, the imputation accuracy of the proposed model was better than those of existing methods in *R*^2^.

**Figure 5:**
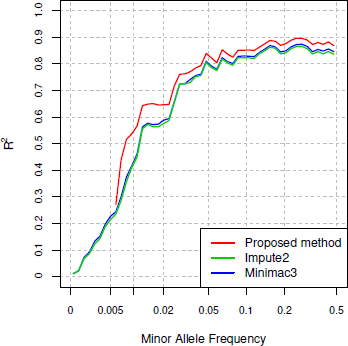
Comparison of *R*^2^ values for the proposed method, Impute2, and Minimac3 for EAS individuals.

## 4 Conclusion

We proposed a genotype imputation method using bidirectional RNN for phased genotype data. Unlike existing imputation methods, the proposed method parameterizes the haplotype information in the reference panel as model parameters in the training step. Since it is difficult to restore genotype data at the individual-level from the trained model parameters, these parameters can be used publicly even when the accessibility of genotype data for training is not permitted publicly. In addition to the simple bidirectional RNN model, we considered the hybrid model comprised of two types of models: one for higher MAF variants and the other for lower MAF variants.

In the performance evaluation using the phased genotype data in the 1000 Genomes Project, we compared settings with the type of RNN cell type, the number of layers, and the number of hidden units in the proposed model, and concluded the model with GRU, 4 layers, and 40 hidden units gave the higher imputation accuracy in terms of *R*^2^ than other settings. We confirmed the effectiveness of the hybrid model from the comparison with models for higher and lower variants. Based on the comparison with existing methods, the imputation performance of the proposed method was comparable or better in specific MAF range and slightly worse in the MAF ≤ 0.05. In the scenario of limited availability of haplotype data for a part of individuals to existing methods, the accuracy of the proposed method was higher than those of the existing methods at least for variants with MAF ≥ 0.005.

## Acknowledgements

This work was supported (in part) by Tohoku Medical Megabank Project from MEXT and Japan Agency for Medical Research and Development, AMED (under Grant Numbers JP19km0105001 and JP19km0105002), and by “Integrative Data Analysis and Data Sharing Promotion for Personalized Prevention and Medicine of Common Diseases” and “Facilitation of R&D Platform for AMED Genome Medicine Support” of Platform Program for Promotion of Genome Medicine from AMED (under Grant Numbers JP19km0405203 and JP19km0405001). This research was also supported by the Center of Innovation Program from Japan Science and Technology Agency, JST. All computational resources were provided by the ToMMo supercomputer system (http://sc.megabank.tohoku.ac.jp/en). We are indebted to all volunteers who participated in this Tohoku Medical Megabank project. We would like to acknowledge all the members associated with this project; the member list is available at the following web site: http://www.megabank.tohoku.ac.jp/english/a171201/.

## S1 Calculation of Dimensionally Reduced Feature Vectors

Let 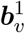 be a binary feature vector of the allele indicated by one for variant *v*. Its *i*th element takes one if the allele of the *i*th haplotype in a reference panel is indicated by one and zero otherwise. We also let 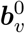 be a binary feature vector of the allele indicated by zero in which the *i*th element takes one if the allele of the *i*th haplotype is indicated by zero and zero otherwise. We apply kernel principal component analysis (PCA) [Schülkopf *et al.*, 1998] to these feature vectors to obtain dimensionally reduced feature vectors. Since the correlation of 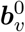 and 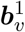 is minus one, we apply kernel PCA only to feature vectors for alleles indicated by one: 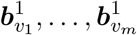, to manage the problem to PCA results caused by highly correlated variables. In order to obtain the dimensionally reduced feature vector of 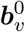, we project 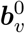 to the space from kernel PCA. We first describe the calculation of kernel PCA briefly, and then derive the projected values of 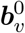 to the space from kernel PCA.

### S1.1 Calculation of kernel PCA

Let *k*(·, ·) and *φ*(·) be a positive definite kernel and the map corresponding to *k* to reproducing kernel Hilbert space (RKHS) 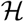, respectively. From the property of RKHS, so called the kernel trick, the inner product 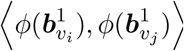 is given by 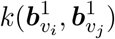. The direction of the first principal component of kernel PCA for 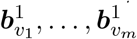 is calculated as follows:

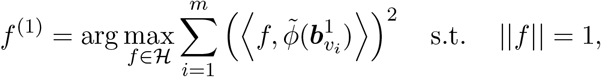

where 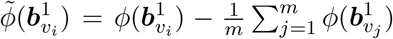 and ∥·∥ indicates the norm in RKHS 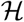. It is sufficient to consider the linear combination of 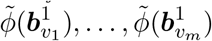 for *f* since directions orthogonal to all the 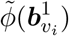 do not contribute to the variance. Thus, the above formula can be rewritten as

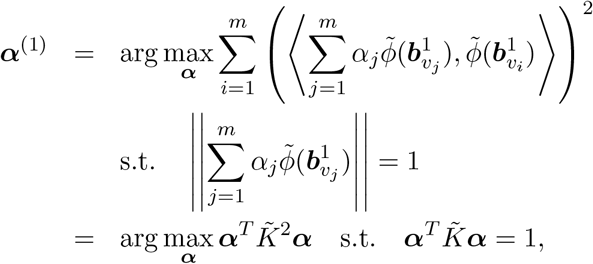

where 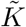 is the centered Gram matrix in which the *i*th row and *j*th column element is given by 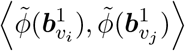. Let *d*_*i*_ and ***u***^(*i*)^ be the *i*th largest eigenvalue of 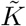 and its corresponding eigenvector, respectively. ***α***^(1)^ is given by 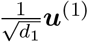, and hence 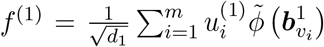, where 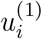 is the *i*th element of ***u***^(1)^. The coefficients for the direction in the ith principal component *f*^(*i*)^ is given by 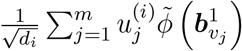 from a similar derivation to *f*^(1)^.

### S1.2 Projection to kernel principal components

Let ***b*** be an original binary feature vector. The *i*th element of the dimensionally reduced feature vector for ***b*** is obtained by its projection to the *i*th kernel principal component, which is also called the *i*th kernel principal component score. The projected value of ***b*** to the *i*th kernel principal component is given as follows:

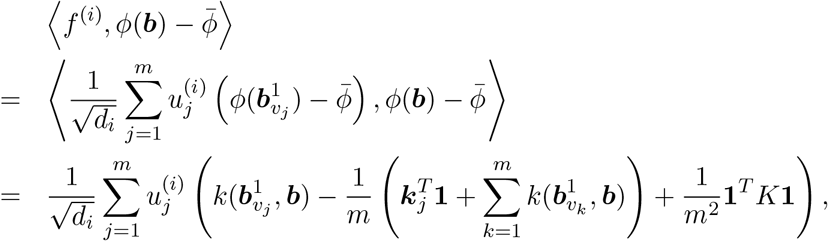

where 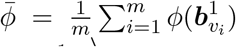, *K* is Gram matrix in which the *i*th row and *j*th column element is 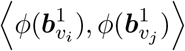, and ***k***_*i*_ is the *i*th column vector of *K*.

